# Deep Proteome Profiling Reveals Signatures of Age and Sex Differences in Paw Skin and Sciatic Nerve of Naïve Mice

**DOI:** 10.1101/2022.07.04.498721

**Authors:** Feng Xian, Julia Regina Sondermann, David Gomez Varela, Manuela Schmidt

**Author notes:** These authors contributed equally (Feng Xian, Julia Regina Sondermann).

## Abstract

The age and sex of studied animals profoundly impact experimental outcomes in animal-based preclinical biomedical research. However, most preclinical studies in mice use a wide-spanning age range from 4 to 14 weeks and do not assess study parameters in male and female mice in parallel. This raises concerns regarding reproducibility and neglects potentially relevant age and sex differences. Furthermore, the molecular setup of tissues in dependence of age and sex is unknown in naïve mice. Here, we employed an optimized quantitative proteomics workflow in order to deeply profile mouse paw skin and sciatic nerve (SCN) – two tissues, which are crucially implicated in nociception and pain as well as diverse diseases induced by inflammation, trauma, and demyelination. Remarkably, we uncovered significant differences when comparing (i) male and female mice, and, in parallel, (ii) adolescent mice (4 weeks) with adult mice (14 weeks). Age was identified as a major discriminator of analyzed samples irrespective of tissue type. Moreover, our analysis enabled us to decipher protein subsets and networks that exhibit differential abundance in dependence on the age and/or sex of mice. Notably, among these were proteins and signaling pathways with known relevance for (patho)physiology, such as homeostasis and epidermal signaling in skin and, in SCN, multiple myelin proteins and regulators of neuronal development. In addition, extensive comparisons with available databases revealed that we quantified approx. 50% of gene products that were implicated in distinct skin diseases and pain, many of which exhibited significant abundance changes in dependence on age and/or sex. Taken together, our study emphasizes the need for accurate age matching and uncovers hitherto unknown sex and age differences at the level of proteins and protein networks. Overall, we provide a unique systems biology proteome resource, which facilitates mechanistic insights into somatosensory and skin biology in dependence on age and sex - a prerequisite for successful preclinical studies in mouse disease models.

**Graphic workflow:** 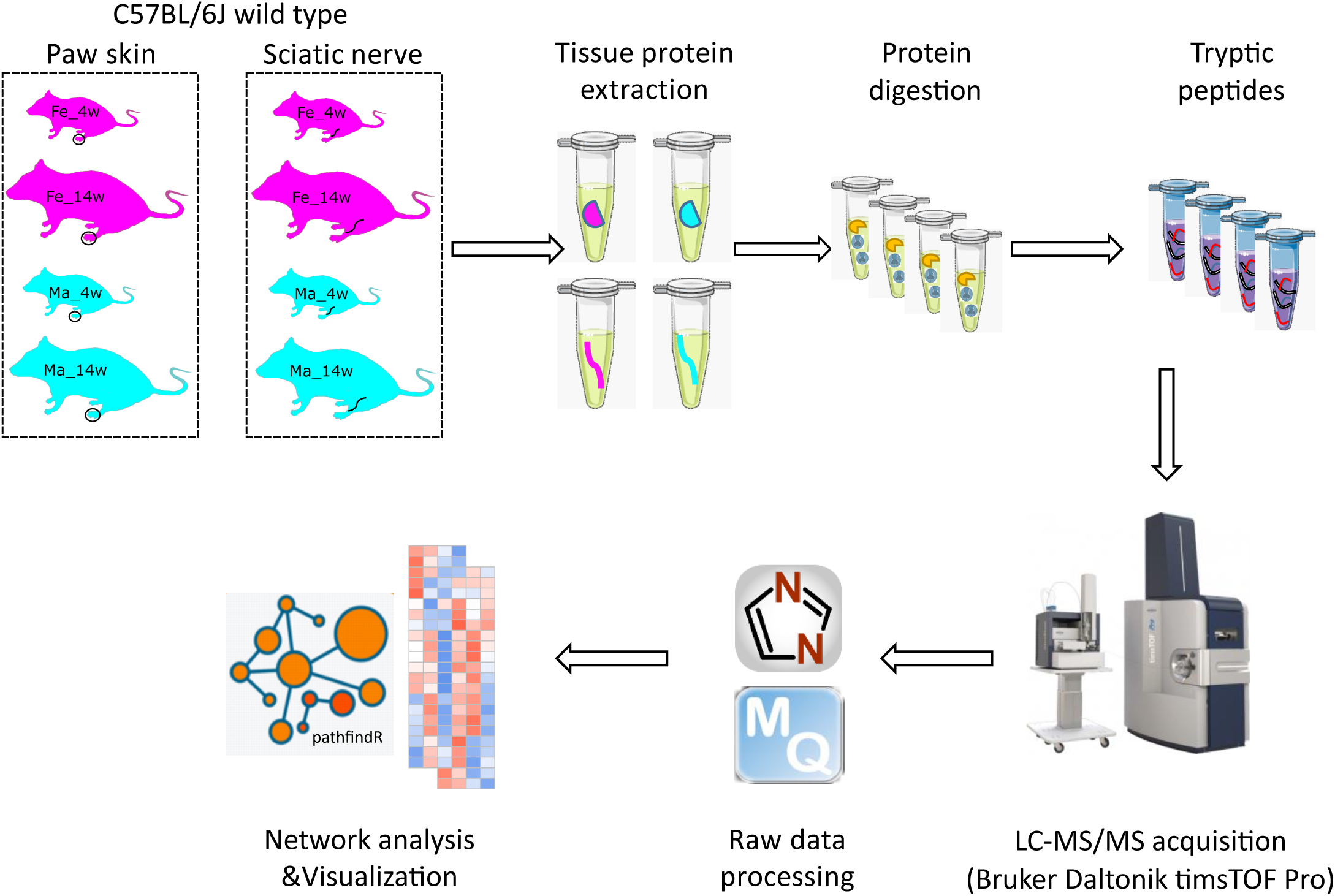

The Figure was partly generated using Servier Medical Art, provided by Servier, licensed under a Creative Commons Attribution 3.0 unported license.

## Introduction

The age and sex of mice are major confounders in preclinical studies, affecting experimental outcomes across scales: from molecular, morphological, and physiological to behavioral parameters (Flórez-Vargas *et al*., 2016; Flurkey *et al*., 2007; Fu *et al*., 2013; Jackson *et al*., 2017). In mice, the first 12 weeks of life are characterized by pronounced changes in terms of growth and development of all organs and systems. This is the reason that Jackson Laboratory (https://www.jax.org) considers the widely used mouse strain C57BL/6J of mature adult physiology only from 12 weeks of age (Flurkey *et al*., 2007). Similarly, the sex of mice needs to be considered when comparing experimental outcomes. Despite recently enforced policies by funding agencies (e.g. NIH policy in 2016) to include animals of both sexes, most preclinical studies still do not perform experiments on male and female rodents in parallel, exhibit gaps in data analysis by sex, and typically pool animals of a wide range of ages (between 4 and 20 weeks) (Flórez-Vargas *et al*., 2016) given time and financial constraints (Garcia-Sifuentes *et al*., 2021; Woitowich *et al*., 2020). These practices may negatively impact reproducibility across studies, increase data variability, conceal differences or generate artefactual results, and, consequently, hamper translationally oriented preclinical research (Flórez-Vargas *et al*., 2016; Jackson *et al*., 2017; Oliva *et al*., 2020).

A prominent example for the enormous diversity of age ranges in publications is studies on rodent (mainly mice and rats) skin and peripheral sensory neurons (e.g. the sciatic nerve, SCN) in the context of somatosensation and pain. Here, it is particularly noteworthy that often different age ranges were used for *in vivo* versus *in vitro* investigations. Mouse behavior experiments assessing paw sensitivity have routinely been performed in mice aged between 6-20 weeks (Hanack *et al*., 2015; Moehring *et al*., 2018; Zheng *et al*., 2019). Studies in cultured peripheral sensory neurons or keratinocytes have used mice aged 4-6 weeks (Hanack *et al*., 2015; Poole *et al*., 2014), 4-8 weeks (Zheng *et al*., 2019), 7-10 weeks (Narayanan *et al*., 2018; Narayanan *et al*., 2016), or 8-16 weeks (Sadler *et al*., 2020). Similarly, myelination of the SCN has been studied biochemically in mice aged 3 weeks (Siems *et al*., 2020), 10 weeks and up to several months (depending on disease severity) (Siems *et al*., 2021), but cultured Schwann cells are generally derived from newborn rats (Siems *et al*., 2020). We have recently discovered a previously unknown age dependence of tactile sensitivity in the back skin and the paw of mice (Michel *et al*., 2020). In particular, 4 weeks old adolescent mice were more sensitive to innocuous tactile stimulation than 12 weeks old adult mice. Interestingly, these observations correlated with similar changes in the activity of the mechanically activated ion channel Piezo2 and age-dependent transcriptome changes in peripheral sensory neurons. Even so, to date, we still lack comprehensive knowledge about the differential molecular set-up of the somatosensory system in dependence on age and sex, in particular on the level of the proteome. This is highly relevant as transcript levels only show limited correspondence with protein levels, which renders the functional interpretation of transcriptome results difficult, in particular under dynamic conditions such as development, maturation, and disease (Liu *et al*., 2016; Schwanhäusser *et al*., 2011; Wang *et al*., 2017). However, in contrast to well-established RNA-seq approaches, deep proteome profiling of complex tissues is still challenging, above all for low abundant and transmembrane proteins. Latest technological advances in mass spectrometry (MS) and data analysis provide new solutions for these challenges (Demichev *et al*., 2020; Meier *et al*., 2020; Meier *et al*., 2018). Here, we thoroughly compared two MS-based quantitative proteomics approaches: commonly used data-dependent acquisition (DDA) paired with parallel accumulation serial fragmentation (DDA-PASEF) (Meier *et al*., 2018) compared to data-independent acquisition DIA-PASEF (Meier *et al*., 2020). The latter has been shown to offer superior performance for deep profiling (Meier *et al*., 2020), yet it has thus far only been applied by highly specialized laboratories given its high demands regarding technology and data analysis.

The goal of this work was to comprehensively catalogue the protein set-up of mouse paw skin and SCN and their changes between adolescent (4 weeks of age) and adult (14 weeks of age) as well as male and female wild-type (WT) C57BL/6J mice. The SCN is affected by a wide variety of motor and sensory neuropathologies induced by inflammation, trauma, and demyelination. Similarly, the skin, as our interface to the outer world, can be impaired by several inflammatory diseases like atopic dermatitis, psoriasis and lupus erythematodes. In addition, both the skin and SCN, are involved in nociception and (chronic) pain. We therefore focused on the potential implication of our data for preclinical research on skin and SCN-related pathologies including pain. Our results decipher hitherto unknown age and sex dependency of diverse proteins and signaling pathways including those with known disease-relevance. Taken together, our dataset is unique as (i) it provides a quantitative protein catalogue of skin and SCN in contrast to the majority of published datasets, which have thus far been performed on the transcriptome level, and (ii) it does so in dependence on age and sex of naïve mice. Given the heterogeneity of mouse age ranges in biomedical studies and the high impact of age and sex on experimental outcomes, our results represent a highly valuable resource to foster future investigations in the context of skin and peripheral nerve (patho)physiology by enhancing reproducibility and unmasking hitherto unknown differences.

## Materials and Methods

### Reagents

All reagents were purchased from Sigma-Aldrich (St. Louis, Missouri) if not mentioned otherwise. Acetonitrile (ACN) and formic acid (FA) were purchased from Fisher Scientific (Hampton, New Hampshire; both FA and ACN were liquid chromatography-mass spectrometry (LC-MS) grade). LC-MS grade water from Sigma was used for all solutions.

### Animals and tissue isolation

In-house bred C57BL/6J mice of both sexes were used. Housing and sacrificing of mice were carried out with approval of the Max Planck Institute for Multidisciplinary Sciences institutional animal care and use committee (IACUC, see Ethics statement). All mice used in this study were group-housed in individually-ventilated cages in a 12 h light/dark cycle in the animal facility of Max Planck Institute for Multidisciplinary Sciences with water and food *ad libitum*. They were sacrificed either at 3-4 or 14-15 weeks. Thus, the experiment consisted of 4 different conditions with each 4 biological replicates (3-4 weeks & female, 3-4 weeks & male, 14-15 weeks & female, and 14-15 weeks & male). After CO_2_-euthanization of mice, the sciatic nerves (SCN) and paw skin were isolated. SCN were rinsed in ice-cold PBS before flash-freezing in liquid nitrogen. For the paw skin, a 4 mm punch biopsy (kai medical, Solingen, Germany, # 48401) of the plantar aspect of the paw was taken, and the dermis and epidermis were separated from underlying tendons/muscle tissue under a microscope. The flash-frozen tissue was stored at - 80°C until further use. For both skin and SCN, tissue from two mice of the same sex and age were pooled together as one biological replicate.

### Protein extraction

For protein extraction, each SCN was cut into 3 pieces with a scalpel on a glass slide and transferred to a protein LoBind tube (Eppendorf, Hamburg, Germany) prefilled with 250 µL lysis buffer (100 mM Tris-HCl, 5% glycerol, 10 mM DTT, 2% SDS) and in presence of 1x complete protease inhibitor cocktail (Roche, Basel, Switzerland). Samples were then sonicated using Bioruptor Pico (Diagenode, Seraing, Belgium) for 15 cycles (30 sec On and 30 sec Off, 4°C) at low frequency. After a short vortex, samples were further incubated at 70°C for 10 min with 1000 rpm agitation. Remaining tissue debris was removed after centrifugation at 10000 g for 5 min, and the supernatant was taken into a new tube. To remove lipids in the tissue lysates, 1250 µL (5x sample volume) of cold acetone was added, and the sample was placed at -20°C for 4 h. With centrifugation at 14000 g for 30 min, acetone was removed, and proteins were collected at the bottom. The protein pellet was further washed with 1.5 mL cold ethanol (80% v/v) followed by 30 min centrifugation at 14000 g. The protein pellet was air-dried for 20 min at room temperature before the addition of 100 µL lysis buffer. A further incubation at 70°C for 10 min with 1000 rpm agitation was performed to solubilize all proteins. Protein concentrations were measured using NanoPhotometer N60 (Implen, München, Deutschland) at 280 nm, and 50 µg protein of each sample was taken for protein reduction (5 mM Dithiothreitol, DTT, 30 min incubation at 60°C) and alkylation (20 mM Iodoacetamide, IAA, 30 min at room temperature in the dark). The remaining IAA in the sample was quenched with addition of 5 mM DTT. Skin biopsies were cut into two pieces and homogenized in 350 µL lysis buffer with the help of a glass dounce. The homogenate was further solubilized by incubation at 70°C for 10 min with 1500 rpm agitation and sonification with the Bioruptor Pico (15 cycles, 30 sec On and 30 sec Off, 4 °C, low frequency). Removal of cell debris and subsequent steps were done as described for the SCN.

### SP3-assisted protein digestion and peptide clean-up

For protein clean-up and digestion, a modified version of the single-pot, solid-phase-enhanced sample preparation (SP3) method from Hughes *et al*. was used (Hughes *et al*., 2019). Briefly, 10 µL of pre-mixed Sera-Mag SpeedBead beads (Cytiva, Marlborough, Massachusetts) were added into 50 µg protein sample. To initiate binding of proteins to the beads, one volume of absolute ethanol was added immediately, followed by incubation on a Thermomixer (Eppendorf) at 24°C for 5 min with 1000 rpm agitation. The supernatant was removed after 2 min resting on a magnetic rack, and the beads were rinsed three times with 500 μL of 80% ethanol. Rinsed beads were reconstituted in 50 μL digestion buffer (50 mM ammonium bicarbonate, pH 8). Protein digestion was performed with 2 μg of either sequencing grade trypsin (SCN samples) or Trypsin/Lys-C (skin samples) for 18 h at 37°C with 950 rpm agitation. After digestion, acetonitrile was added to each sample to a final concentration of 95%. Mixtures were incubated for 8 min at room temperature and then placed on a magnetic rack for 2 min. The supernatant was discarded, and the beads were rinsed with 900 μL of 100% ACN. Rinsed beads were reconstituted either in 40 μL (SCN samples) or 20 μL (skin samples) LC-MS grade water to elute the peptides. Peptide concentration was measured in duplicate using NanoPhotometer N60 (Implen, München, Deutschland) at 205 nm. Peptide samples were acidified with FA to a final concentration of 0.1% and stored at -20°C until LC-MS/MS analysis.

### LC-MS/MS

Nanoflow reversed-phase liquid chromatography (Nano-RPLC) was performed on a NanoElute system (Bruker Daltonik, Bremen, Germany). 250 ng of peptides were separated with a 130 min gradient on a 25 cm x 75 µm column packed with 1.6 µm C18 particles (IonOpticks, Fitzroy, Australia). Mobile solvent A consisted of 2% ACN, 98% water, 0.1% FA and mobile phase B of 100%, 0.1% FA. The flow rate was set to 400 nL/min for the first 2 min and the last 9 min of the gradient, while the rest of the gradient was set to 250 nL/min. The mobile phase B was linearly increased from 0 to 20% from 3 min to 110 min, flowed by a linear increase to 35% within 10 min and a steep increase to 85% in 0.5 min. Then a flow rate of 400 nL/min at 85% was maintained for 9 min to elute all hydrophobic peptides. NanoElute LC was coupled with a hybrid TIMS quadrupole TOF mass spectrometer (timsTOF Pro, Bruker Daltonik) via a CaptiveSpray ion source. Each sample was analyzed in both data-independent acquisition (DIA) and data-dependent acquisition (DDA) modes coupled with parallel accumulation serial fragmentation (PASEF) one after another in duplicate. The TIMS analyzer was operated in a 100% duty cycle with equal accumulation and ramp times of 100 ms each. Specifically, in DDA-PASEF mode (Meier *et al*., 2018), 10 PASEF scans were set per acquisition cycle with ion mobility range (1/k0) from 0.6 to 1.6, and singly charged precursors were excluded. Dynamic exclusion was applied to precursors that reached a target intensity of 17500 for 0.4 min. Ions with m/z between 100 and 1700 were recorded in the mass spectrum. In DIA-PASEF mode, precursors with m/z between 400 and 1200 were defined in 16 scans containing 32 ion mobility steps with an isolation window of 26 Th in each step with 1 Da overlapping for neighboring windows. The acquisition time of each DIA-PASEF scan was set to 100 ms, which led to a total cycle time of around 1.8 sec (Meier *et al*., 2020). In both DDA and DIA-PASEF modes, the collision energy was ramped linearly from 59 eV at 1/k0 = 1.6 to 20 eV at 1/k0 = 0.6.

### DDA-PASEF data processing

All DDA data were analyzed with MaxQuant (version 1.6.17.0) and searched with Andromeda against *Mus musculus* database from Uniprot containing 17070 protein entries (downloaded on 2021-07-08). The minimal peptide length was set to 6 amino acids, and a maximum of 3 missed cleavages were allowed. The search included variable modifications of methionine oxidation and N-terminal acetylation, deamidation (N and Q) and fixed modification of carbamidomethyl on cysteine, and a maximum of 3 modifications per peptide were allowed. The “Match between run” function was checked within 0.5 min retention time window and 0.05 ion mobility window. Mass tolerance for peptide precursor and fragments were set as 10 ppm and 20 ppm, respectively. The FDR was set to 0.01 at precursor level and protein level. Label-free quantification algorithm was used to quantify identified proteins with a minimum 1 razor and unique peptide. The rest of the parameters were kept as default. Proteus, an R package (https://github.com/bartongroup/Proteus), was used for downstream analysis of MaxQuant output (Gierlinski *et al*., 2018).

### DIA-PASEF data processing

DIA-NN (Demichev *et al*., 2020) was used to process DIA-PASEF data in library-free mode with the same *Mus musculus* proteome database to generate the predicted spectrum library. Trypsin/P was used for *in silico* digestion with an allowance of maximum 3 missed cleavages. A deep learning-based method was used to predict theoretical peptide spectra along with its retention time and ion mobility. Variable modifications on peptides were set to N-term methionine excision, methionine oxidation and N-term acetylation, while carbamidomethylation on cysteine was a fixed modification. A maximum number of variable modifications on a peptide was set to 3. Peptide length for the search ranged from 5 to 52 amino acids. Aligned with the DIA-PASEF acquisition method, m/z ranges were specified as 400 to 1200 for precursors and 100 to 1700 for fragment ions. Both MS1 and MS2 mass accuracy were set to 10 ppm as recommended. Unique genes were used as protein inference in grouping. RT-dependent cross-run normalization and Robust LC (high accuracy) options were selected for quantification. The main report from the DIA-NN search was further processed with the R package, DIA-NN (https://github.com/vdemichev/diann-rpackage), to extract the MaxLFQ (Cox *et al*., 2014) quantitative intensity of gene groups for all identified protein groups with q-value < 0.01 as criteria at precursor and gene group levels.

### Visualization of proteomic data

Pearson’s correlation plots were created with corrplot package (https://github.com/taiyun/corrplot). Venn diagrams were plotted using VennDiagram (https://CRAN.R-project.org/package=VennDiagram), and area-proportional Venn diagrams were created with eulerr package (https://github.com/jolars/eulerr). All box plots and bar plots used to visualize the proteomic data were created using ggplot2 package (https://github.com/tidyverse/ggplot2). In addition, the package ggpubr (https://rpkgs.datanovia.com/ggpubr/) was used for significance tests in comparisons within box plots. Principal component analysis was performed with the factoextra package (https://rpkgs.datanovia.com/factoextra/index.html). Color-coded tables were prepared in Microsoft Excel 2019.

### Differential expression analysis and protein function enrichment

Two samples (Skin_Ma4w-4 and SCN_Fe4w-1) were excluded from the following analysis as we noticed that these two samples were outliers in principal component analysis (PCA) analysis, and protein content of these two samples was significantly lower than in corresponding biological replicates. Quantitative data was imported into the R package, ProTIGY (https://github.com/broadinstitute/protigy) and log2-transformed. Quantified proteins presented in less than 25% of all replicates were filtered out, and 7323 protein groups from paw skin and 7144 protein groups from SCN were submitted to differential expression analysis. Two-sample moderated T-test was used to test for statistical significance test of individual contrasts. Age-dependent comparisons (Fe = female; Ma = male): Skin_Fe4w versus Skin_Fe14w, Skin_Ma4w versus SkinMa14w, SCN_Fe4w versus SCN_Fe14w, and SCN_Ma4w versus SCN_Ma14w. Sex-dependent comparisons: Skin_Ma4w versus Skin_Fe4w, Skin_Ma14w versus Skin_Fe14w, SCN_Ma4w versus SCN_Fe4w, and SCN_Ma4w versus SCN_Fe4w. Proteins with the adjusted (Benjamini and Hochberg, for multiple testing) p-value ≤ 0.05 (hereafter referred to as the q-value) and the absolute log2 fold change (FC) ≥ 0.585 were considered as differentially expressed proteins (DEPs) in each contrast. Gene Ontology Biological Process (GO-BP) enrichment and visualization of DEPs was performed using the pathfindR package (https://github.com/egeulgen/pathfindR) in R environment (Ulgen *et al*., 2019). A minimal of 4 DEPs and adjusted p-value ≤ 0.05 (Bonferroni) were applied for significantly enriched pathways.

## Results

### DIA-PASEF allows deep and reproducible proteome profiling of mouse paw skin and sciatic nerve

In this study, we analyzed 16 biological replicates of paw skin and sciatic nerve (SCN) samples to compare the proteome between (i) two age groups, i.e. 4 weeks old adolescent mice and 14 weeks old adult mice, and (ii) males and females. To enable and optimize truly deep proteome profiling, we compared two label-free quantification strategies of mass spectrometry (MS)-based quantitative proteomics. In particular, data-dependent acquisition parallel accumulation serial fragmentation (DDA-PASEF), and data-independent acquisition (DIA)-PASEF. For each sample, we analyzed technical duplicates using a timsTOF Pro mass spectrometer (Bruker Daltonik). DDA or DDA-PASEF have been the methods of choice for most proteomics studies published so far (Aballo *et al*., 2021; Aebersold *et al*., 2016; Meyer, 2021). However, recent advances highlighted the superior performance of DIA-PASEF methods (Brunner *et al*., 2022; Meier *et al*., 2021), which we tested in our study side-by-side. Given the long acquisition time of approx. 20 days for all samples and replicates, we constantly monitored the performance of our MS-setup in DIA-PASEF mode by using pooled skin peptides and SCN peptides as quality controls. Pearson’s correlation coefficients were calculated for all quality control runs (Figure 1A, B). The average correlations of quality controls were 0.98 and 0.99 for pooled skin and SCN samples, respectively, indicating highly consistent stability of the instrument-setup. Usually, DIA data is searched against a peptide library constructed from data obtained via DDA of the same sample, therefore, only proteins present in the library can be identified and quantified (Ludwig *et al*., 2018). In contrast, DIA-NN, a recently developed program based on deep neural networks, extensively advanced DIA workflows with a library-free database search mode (Demichev *et al*., 2020). Thus, we compared DDA-PASEF data subjected to a standard MaxQuant search (Cox *et al*., 2014) with DIA-PASEF data subjected to DIA-NN library-free search. As shown in Figure 1C-D, protein identifications from DDA-PASEF were highly covered by DIA-PASEF experiments, and DIA-PASEF detected additional 4317 and 3926 protein groups in skin and SCN, respectively (Figure 1-source data 1, 2). Besides comparing protein identifications (protein IDs: for the remainder of this manuscript we will refer to protein groups as protein IDs for the sake of simplicity), we also compared both acquisition modes in respect to reproducibility at the quantitative level. Notably, we observed smaller coefficients of variation across all DIA-PASEF runs (Figure 1E, F), indicating improved reproducibility over DDA-PASEF. Taken together, DIA-PASEF exhibited superior performance and was therefore chosen for further analysis of skin and SCN samples.

**Figure 1:**
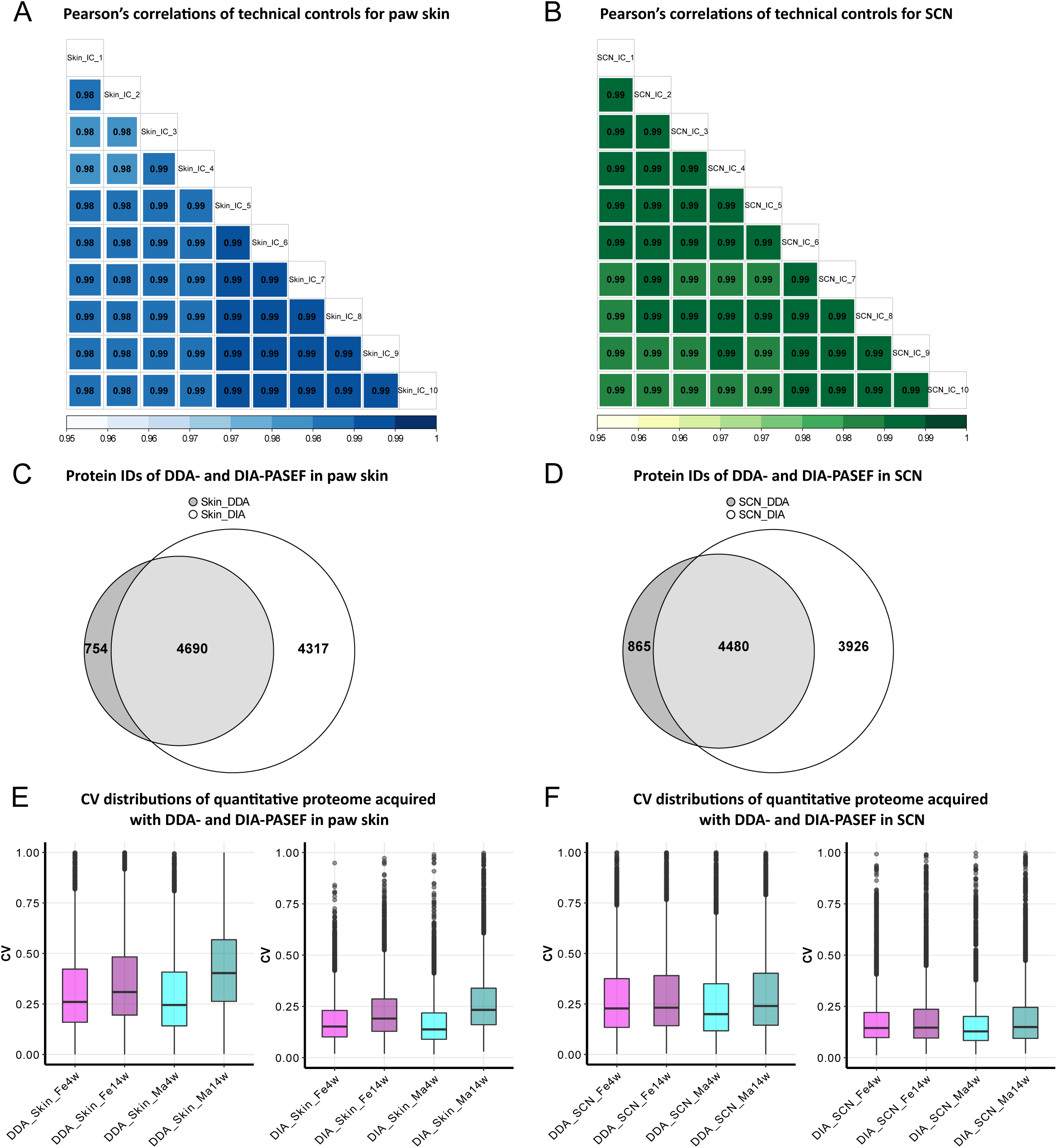
DIA-PASEF acquisition followed by DIA-NN analysis outperforms DDA-PASEF acquisition in deep proteome profiling of paw skin and SCN of naïve mice. A-B: Pearson’s correlations of technical controls of paw skin (blue) and SCN (green) acquired over 20 days on a timsTOF Pro. C-D: Comparisons of quantified protein groups (protein IDs) using DDA- and DIA-PASEF workflows in paw skin (C) and SCN (D). E-F: Coefficient of variation (CV) distributions of quantitative proteomes using DDA- and DIA-PASEF in paw skin (E) and SCN (F) of 4 weeks and 14 weeks old males (cyan) and females (magenta).

### Age-dependent protein abundance changes in mouse paw skin and SCN

In paw skin, we quantified over 8600 protein IDs across experimental groups (Figure 2A, Figure 1-source data 1). Comparing this proteome dataset with the most comprehensive (human) skin proteome dataset (Dyring-Andersen *et al*., 2020) published so far, our skin proteome covered over 70% (Figure 2B). Importantly, in our study we analyzed whole skin lysates without pre-analytical sample fractionation (e.g. into different skin layers) (Dyring-Andersen *et al*., 2020). Note that the previously published skin proteome was obtained from human hairy skin, while we analyzed mouse glabrous skin known to exhibit several differences in skin structure (Gudjonsson *et al*., 2007). Nonetheless, we identified all 50 known keratins and 19 collagens. In addition to structural proteins, we also quantified 13 members of the interleukin (IL) family and 11 of the S100 family (Figure 2E), known to play essential roles in the context of inflammation and infection (Kozlyuk *et al*., 2019; Velazquez-Salinas *et al*., 2019). Their detection across all skin samples with only a few missing values (note that we did not impute any data, please see methods for details), further validates the high performance and reproducibility of our optimized proteomic workflow. In SCN, over 8400 protein IDs were quantified across experimental groups (Figure 2C, Figure 1-source data 2). SCN harbors myelinated axons, which are closely associated with glia cells such as Schwann cells. Remarkably, the myelin proteome was nearly completely covered in our SCN data (94%; 1014/1077 described myelin proteins (Siems *et al*., 2020); Figure 2D), without *a priori* myelin enrichment as required in previous studies (Siems *et al*., 2020). Among the 63 proteins of the myelin proteome, which were not covered in our dataset, were mainly ATP synthases, Histones, and Septins (Figure 2-source data 1). Another indication as to the depth and high performance of our workflow is the fact, that we robustly quantified multiple ion channels across SCN samples (Figure 2F, Figure 1-source data 2), such as Trpv1 and several voltage-gated sodium channels (e.g. Scn8a, Scn9a, Scn11a) – again without requiring pre-analytical membrane preparations. These ion channel identifications further corroborate the high quality of our approach as ion channels are usually expressed at low abundance and are notoriously difficult to be detected by MS given their pronounced hydrophobicity (Samways, 2014).

**Figure 2:**
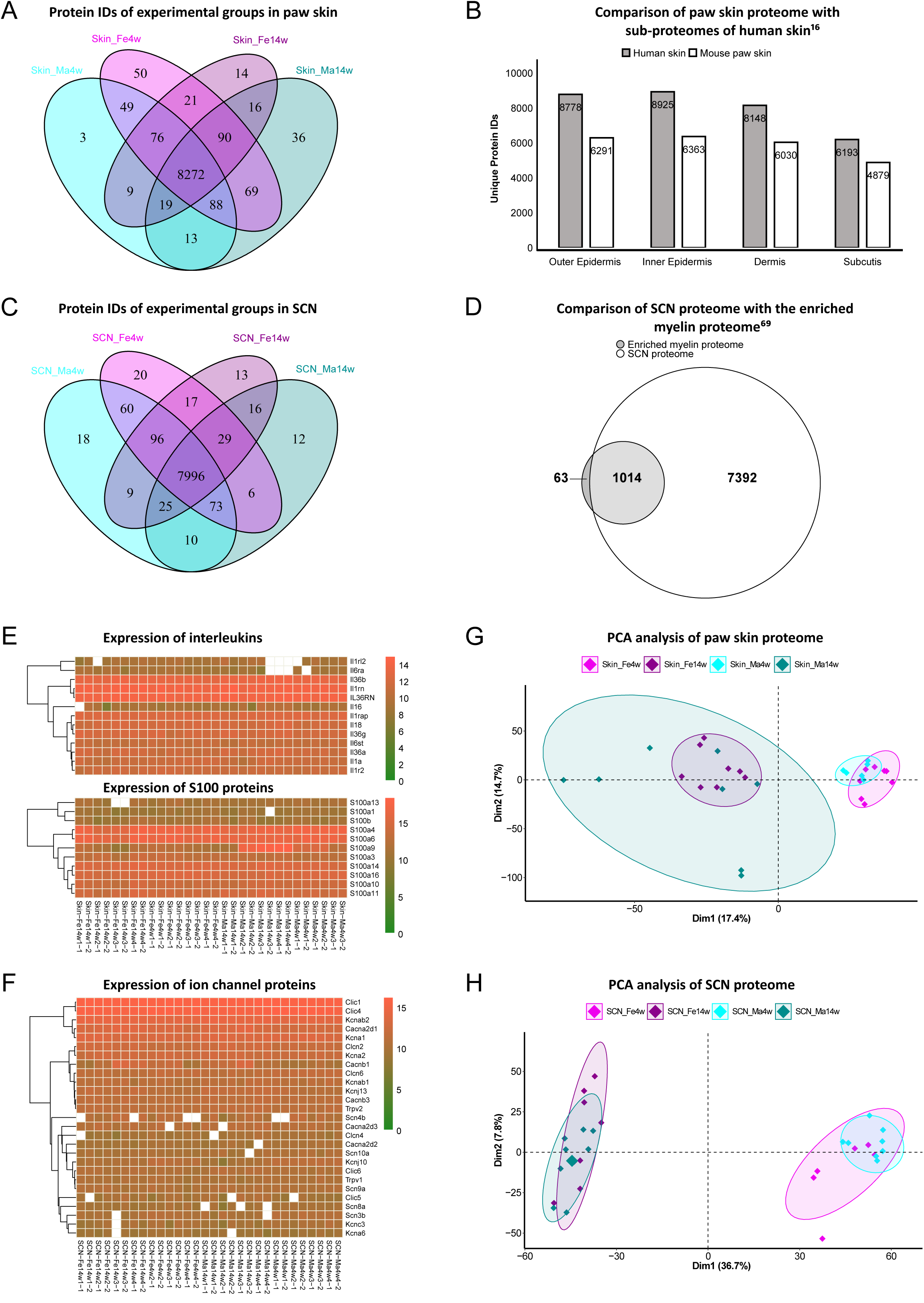
Age and sex differences in proteomes of paw skin and SCN. A: Venn diagram shows unique and shared protein IDs across age and sex groups of paw skin. B: Comparison of the quantified paw skin proteome with previously reported sub-proteomes of human skin (Dyring-Andersen *et al*., 2020) indicates high coverage of our proteome data. C: Venn diagram shows unique and shared protein IDs across age and sex groups of SCN. D: Our SCN proteome dataset harbors 1014 myelin proteins (94%) of the previously reported myelin proteome (Siems *et al*., 2020). E: Heatmaps show the expression of interleukin and S100 protein families across all paw skin samples. F: Heatmaps show the expression of ion channel proteins quantified across all SCN samples. Color legends are coded based on log2-transformed protein intensities. G-H: Principal component analysis (PCA) reveals age as a prominent variable in paw skin and SCN tissues.

We employed principal component analysis (PCA) to visualize proteome similarities and differences across age and sex groups. Importantly, we only considered those proteins that were robustly quantified in all samples (according to all our quality criteria, please see methods for details), resulting in 6086 protein IDs in the skin and 6075 protein IDs in SCN (Figure 2G, H; Figure 1-source data 1, 2). Age groups were clearly segregated in both skin and SCN samples by the first and second component, indicating that age is a prominent discriminator in our study and associated differences can be tackled by whole proteome analysis. Furthermore, to elucidate changes in abundance profiles across all experimental groups, fuzzy C-means clustering analysis was performed based on the average quantitative intensity of each protein ID (Figure 2–figure supplement 1). Among the 9 clusters generated, most of the proteins showed strong age patterns, such as Clusters 2 and 5 in skin, and Clusters 4, 6 and 7 in SCN. On the contrary, several proteins exhibited different expression trends in age/sex groups. For instance, most proteins in Cluster 6 of the skin proteome showed minor age-dependent changes in females, while their abundance was notably increased in 14 weeks males compared to 4 weeks males (Figure 2–figure supplement 1). Similar sex-specific changes were also observed in SCN represented by Cluster 3. Taken together, the clustering analysis of the paw skin and SCN proteome reveals thus far unknown expression patterns dependent on the biological variables age and sex, i.e. sex-specific and overlapping age dependency, which may affect mouse (patho)physiology.

### Diverse biological pathways exhibit age dependency in paw skin

To explore this age dependency further, we applied a fold change (FC) cutoff (absolute log2 FC ≥ 0.585) in addition to q-value ≤ 0.05 and found 588 and 503 highly differentially expressed proteins (DEPs) in female and male skin datasets (Figure 1-source data 1). As shown in Figure 3A, 211 DEPs were shared in between sexes, while 377 and 292 DEPs were unique for female and male skin, respectively (Figure 1-source data 1). Importantly, among the 211 common DEPs, we noticed a set of DEPs with opposite age-dependent expression when comparing male and female mice. Most of these proteins had higher expression levels in 4 weeks female mice, while their expression was lower in 4 weeks male mice (Figure 3B). Gene Ontology Biological Process (GO-BP) analysis of 211 common DEPs resulted in 5 significantly enriched pathways (criteria: at least 4 DEPs/pathway, Bonferroni adjusted p-value ≤ 0.05). DEPs annotated to enriched pathways were mapped back to the quantitative proteomic data, and the agglomerated scores of the pathways are visualized in Figure 3C, revealing a marked age-dependent pattern. As expected, most of the enriched pathways were related to skin development. For instance, ‘protein hydroxylation’ and ‘collagen fibril organization’ were both more “active” in 4 weeks skin than at 14 weeks. Specifically, several proline/serine hydroxylases (e.g. P3h4, P4ha2, P3h1, P4ha1) were highly expressed in 4 weeks skin together with members of collagens (Figure 3–figure supplement 1A). These two pathways were reported to be essential for skin stability during development (Rappu *et al*., 2019). On the contrary, the ‘cornification pathway’ appeared to be more “active” in 14 weeks mice as expected, given its role in skin maturation. Performing GO-BP enrichment on unique DEPs from age-dependent comparisons in female and male mice (Figure 3–figure supplement 1B, C) revealed interesting biological insights into sex-dependent differences. In male mice, pathways of ‘sodium ion transmembrane transport’, ‘substrate adhesion dependent cell spreading’, and ‘regulation of cell shape’ were significantly enriched at 4 weeks, and ‘response to mechanical stimulus’ at 14 weeks, respectively (Figure 3D). These pathways were inter-connected via a few proteins, for instance, Polycystin-2 (Pkd2) and Dystrophin (Dmd), both of which being more abundant in 4 weeks male skin. Pkd2 functions as a component of a calcium-permeable ion channel (Kim *et al*., 2016), and can form mechano- and thermo-sensitive channels together with Transient receptor potential cation channel subfamily V member 4 (Trpv4) (Kottgen *et al*., 2008). Pkd2 might also be involved in cell spreading: The increase of traction forces between cell and substrate may cause higher cortical tension, which in turn may open mechanosensitive channels (Xie *et al*., 2018). Our results may be indicative of a generalized higher activity of ion transport systems across the membrane in younger male mice, which could potentially influence the sensitivity to stimuli applied to the skin. Indeed, we previously reported increased tactile sensitivity in 4 weeks mice compared to adults (Michel *et al*., 2020) and future experiments will need to address whether DEPs identified here might contribute to this phenomenon. Note, however, that we did not observe any other ion channel to be differentially expressed by age in our male skin datasets (Figure 1-source data 1). In female skin, most proteins annotated to interconnected pathways of ‘regulation of action potential’, ‘response to nerve growth factor’, and ‘response to acid chemical’ were significantly enriched at 4 weeks compared to 14 weeks (Figure 3E). Similarly, several mitochondrial annotated pathways and regulation of reactive oxygen species were enriched in 4 weeks female skin (Figure 3–figure supplement 1B), all of which have been reported to contribute to skin homeostasis (Hamanaka *et al*., 2013; Sreedhar *et al*., 2020).

**Figure 3:**
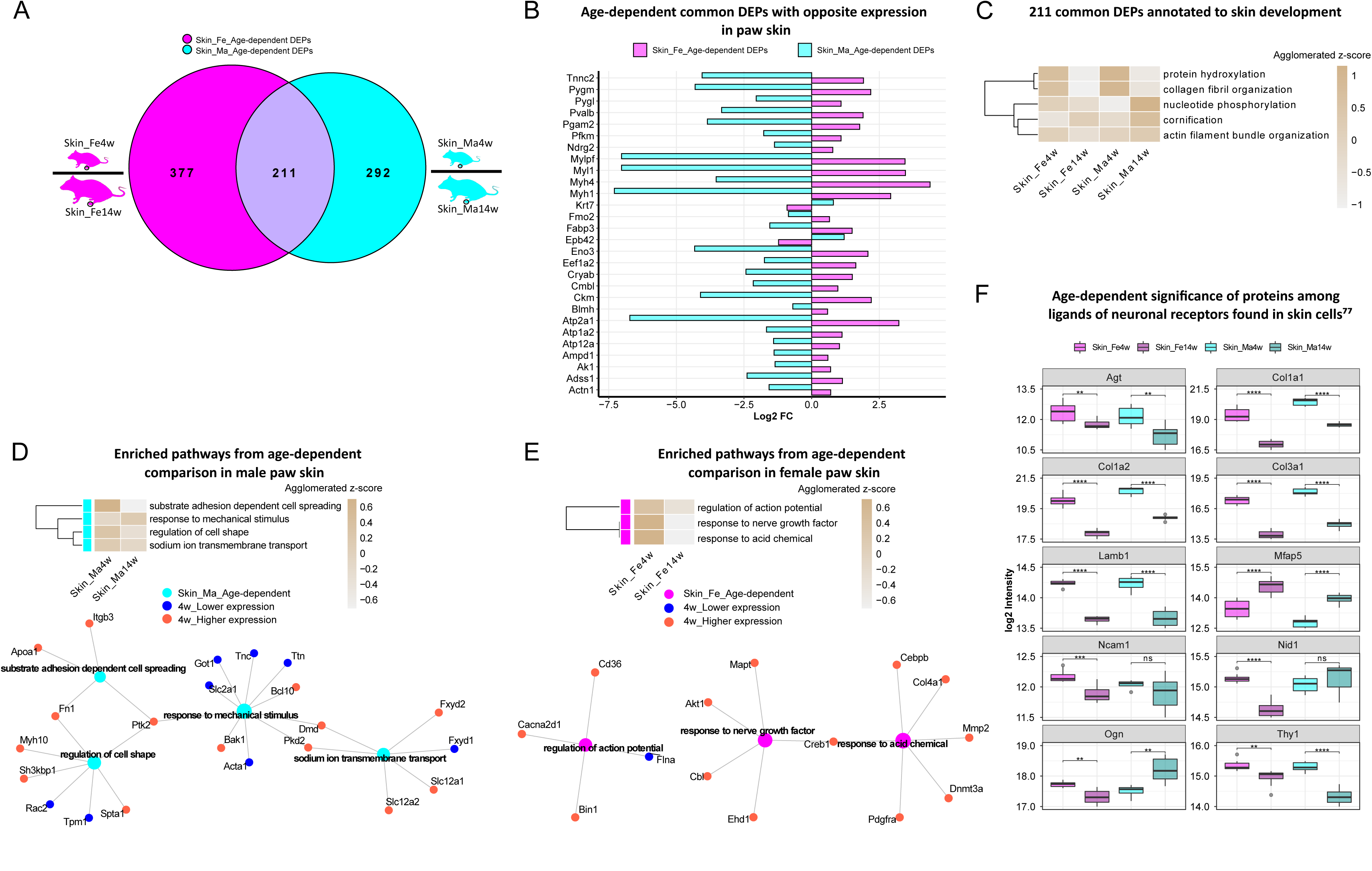
Differential expression analysis of paw skin samples reveals diverse age-dependent biological pathways. A: Venn diagram illustrates unique and shared differentially expressed proteins (DEPs; criteria: q-value ≤ 0.05, absolute log2 FC ≥ 0.585) from age-dependent comparisons in female (magenta) and male (cyan) paw skin. B: Bar plot compares a set of common DEPs with opposite expression when comparing 4 weeks with 14 weeks females and males. C: 211 common DEPs (please see A) are annotated to pathways related to skin development. The agglomerated z-score of each pathway is visualized in the heatmap. D-E: Enriched interconnected pathways from age-dependent comparison in male and female mice. Red: higher expression at 4 weeks; blue: lower expression at 4 weeks. F: Eight ligands of neuronal receptors found in skin cells (Wangzhou *et al*., 2021) are significantly regulated by age. Significance levels are indicated as: ns (q-value > 0.05), * (q-value ≤ 0.05), ** (q-value ≤ 0.01), *** (q-value ≤ 0.001), and **** (q-value ≤ 0.0001).

Keratinocytes are by far the most abundant cell types in skin, followed by fibroblasts, endothelial cells, melanocytes, and subsets of resident innate and adaptive immune cells. In addition, sparsely distributed sensory nerve endings in the skin play significant roles for all aspects of somatosensation, including the detection of different physical stimuli, may they be innocuous or noxious. However, this cellular diversity cannot be separated on the experimental level when analyzing complex tissue lysates as in our study. In contrast, we assessed the depth of our profiling workflow across different cell types indirectly by applying a recently published resource on ligand-receptor interactions in 42 cell types, including sensory neurons of mouse dorsal root ganglia (DRG) (Wangzhou *et al*., 2021). We extracted ligand-receptor interactions found across skin cell types (Wangzhou *et al*., 2021) for comparison with our skin dataset. In total, 52 ligands of DRG were present in our skin dataset (Figure 3-source data 1), and 8 of them were significantly regulated (q-value ≤ 0.05) when comparing 4 weeks to 14 weeks mice of both sexes (Figure 3F). For example, the ligand Lamb1 was found to be more abundant in 4 weeks skin. Lamb1 was reported to serve as an anchor point for end feet of radial glial cells and as a physical barrier to migrating neurons (Radmanesh *et al*., 2013). Three receptors of Lamb1, low-density lipoprotein receptor-related protein 1 (Lrp1), C-type mannose receptor 2 (Mrc2), and Suppressor of tumorigenicity 14 protein homolog (St14), were also identified in our dataset (Figure 1-source data 1). Interestingly, Lrp1 and Mrc2 showed age-dependent statistical significance with higher expression at 4 weeks of age. Further, Thy1, a membrane glycoprotein more abundant in 4 weeks in male and female mice (Figure 3F), was shown to be involved in cell-cell or cell-ligand interactions during synaptogenesis (Zwerner *et al*., 1977), suggesting a hitherto unknown and possible role in skin maturation. Apart from the functional importance of those ligands during skin development, our results generally raise awareness of the pronounced age dependency of protein expression in naïve mice, which should be carefully considered when pooling wide-ranging age groups in mouse studies.

### Prominent age dependency of immune pathways and myelin proteins in SCN

In the SCN proteome, we observed similar age dependency as in paw skin. Differential expression analysis uncovered 1128 DEPs and 1595 DEPs in age-dependent comparisons of female and male SCN (Figure 4A, Figure 1-source data 2), accounting for almost one quarter of the SCN proteome (Figure 2C). Pathway enrichment for 803 common DEPs (Figure 1-source data 2), and age-enriched DEPs (for 4 weeks and 14 weeks, respectively) is given in Figure 4–figure supplement 1, spanning diverse categories from metabolic processes and translation to inflammatory and immune signaling. For example, among common DEPs, ‘response to epidermal growth factor’ and ‘glial cell differentiation’ pathways had a higher z-score in 4 weeks mice of both sexes, while pathways related to ‘neuron survival’ and ‘axonal transport’ were more pronounced in 14 weeks mice of both sexes (Figure 4B). Again, these processes appear to be interconnected via distinct DEPs (Figure 4B), suggesting synergies during development. For instance, Superoxide dismutase (Sod1) was found to be less expressed in 4 weeks mice and represents a connecting hub of 3 pathways related to nervous system function (Figure 4B) in line with its implication in amyotrophic lateral sclerosis (ALS) (Pansarasa *et al*., 2018). Remarkably, 177 DEPs from age-dependent comparisons were associated with the ‘synapse’ as revealed by querying SynGO, a public reference for synapse research (Koopmans *et al*., 2019) (Figure 4-source data 1).

**Figure 4:**
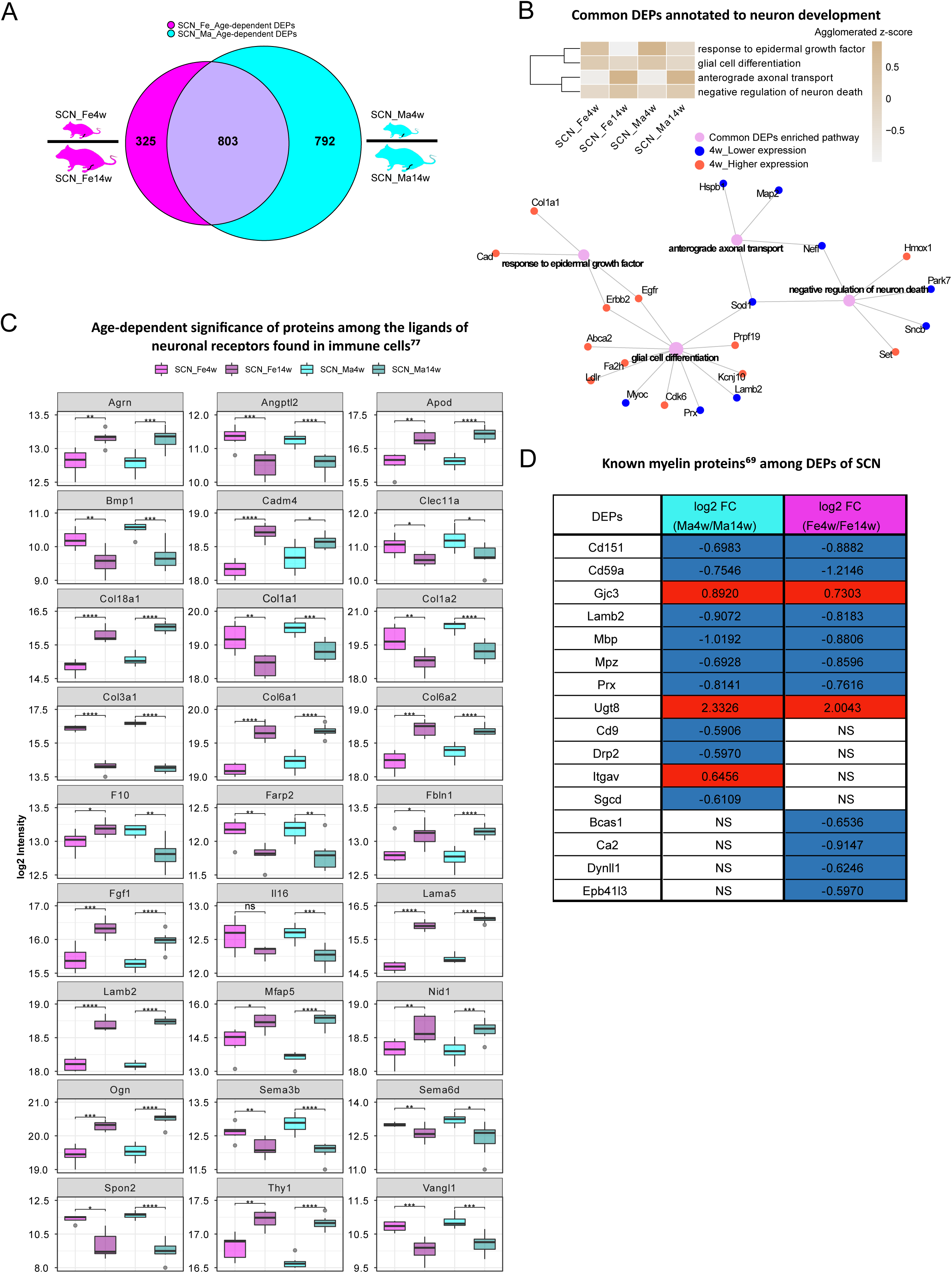
Age-dependent differential expression analysis in SCN samples. A: Venn diagram illustrates unique and shared DEPs (criteria: q-value ≤ 0.05, absolute log2 FC ≥ 0.585) from age-dependent comparisons in female (magenta) and male (cyan) SCN. B: Common DEPs are annotated to pathways related to neuron growth. The agglomerated z-score of each pathway was visualized in the heatmap. Red: proteins more abundant at 4 weeks; blue: proteins less expressed at 4 weeks. C: Twenty-seven ligands of neuronal receptors found in immune cells (Wangzhou *et al*., 2021) are significantly regulated by age. D: Log2 FC of previously reported myelin proteins (Siems *et al*., 2020), which are significantly regulated by age. Red: higher expression at 4 weeks; blue: lower expression at 4 weeks; NS: not significantly regulated. Significance levels are indicated as: ns (q-value > 0.05), * (q-value ≤ 0.05), ** (q-value ≤ 0.01), *** (q-value ≤ 0.001), and **** (q-value ≤ 0.0001).

Diverse immune cell types are known to be presented in the SCN, where they contribute to nerve health, damage, and repair as well as to sensory phenomena of pain (Kalinski *et al*., 2020). Thus, we cross-referenced our data to aforementioned ligand-receptor database (Wangzhou *et al*., 2021) and searched our SCN dataset for ligands of neuronal receptors known to be expressed in immune cells. Among the 57 immune cell ligands of neuronal receptors quantified in the SCN proteome, 27 showed age-dependent abundance changes such as Agrin (Agrn), Pro-interleukin IL-16 (Il16), and several collagens (Figure 4C, Figure 4-source data 2). Given their age-dependent abundance differences already in naïve mice, our results caution to adequately pool age groups when assessing immune signaling in mouse disease models in future studies as data might get skewed by underlying – and thus far unknown – age differences.

We also assessed ligands of neuron receptors (Wangzhou *et al*., 2021) found in glial cells and vice versa, given their utmost importance for SCN (patho)physiology. We found 87 glial cell ligands of neuron receptors and 71 neuron ligands of glial cell receptors in our SCN proteome, and, more interestingly, about one third of ligand-receptor pairs showed strong age dependency (Figure 4-source data 3, 4). For example, Limbic system-associated membrane protein (Lsamp), a glial cell ligand mediating selective neuronal growth and axon targeting (Sanz *et al*., 2017), and two of its neuronal receptors, Netrin-G1 (Ntng1) and Thy-1 membrane glycoprotein (Thy1). While Lsamp exhibited higher expression in 4 weeks SCN than in 14 weeks SCN, its two receptors showed the opposite (Figure 4-source data 4). These data may suggest a homeostatic mechanism specifically in young SCN to counterbalance higher Lsamp abundance on its receptor level – an intriguing hypothesis given that Lsamp has been reported to suppress neuronal outgrowth of DRG (Sanz *et al*., 2017).

In this respect, it is noteworthy that 16/90 known myelin proteins (Siems *et al*., 2020) within the myelin proteome (Figure 2D) were differentially regulated by age, including highly abundant structural myelin proteins such as Myelin basic protein (Mbp) and Myelin protein P0 (Mpz) (Figure 4D). In line with previous reports in mice and zebrafish (Siems *et al*., 2021), Mbp and Mpz were significantly enriched in both male and female 14 weeks SCN, reflecting myelin assembly and axonal development with age. Similarly, CD59A glycoprotein (Cd59), a sparsely expressed myelin protein associated with protection against complement-mediated lysis (Zeis *et al*., 2016), was more abundant at adult age (Figure 4D) as described previously in mouse brains (Siems *et al*., 2021). This congruency with published data on both, high and low abundant myelin proteins, validates our datasets and highlights their quality and depth of profiling. Note, however, that many myelin DEPs appear to be specific for male or female SCN in dependence on age – a fact, which has not been investigated in previous studies which focused on male mice only (Siems *et al*., 2021). For example, the CD9 antigen (Cd9) is implicated in immunomodulation within myelin (Zeis *et al*., 2016), and our datasets show a higher abundance in 14 weeks males compared to 4 weeks males (Figure 4D). Interestingly, female mice did not exhibit this age-dependent regulation of Cd9 (Figure 4D). Further investigation of these age- and sex-dependent changes will likely help to better understand the molecular set-up of myelin and, importantly, associated pathologies.

### Sexual dimorphism in paw skin and SCN proteomes

We then turned to specifically looking at sex differences in our datasets. While several studies have addressed this issue in animal models, most previous reports relied on investigating differences of transcript abundance in paw skin and nerve tissues (Mecklenburg *et al*., 2020; Ray *et al*., 2019). In our proteome dataset we observed sex-dependent changes in skin and SCN with 292 DEPs and 75 DEPs, respectively (Figure 5A, B; Figure 1-source data 1, 2). Yet, less so when compared to aforementioned prominent age dependency (Figures 3A, 4A). Also, here it is noteworthy that sex-differences were dependent on age. For example, they were less pronounced in 4 weeks compared to 14 weeks skin and SCN (Figure 5A, B, Venn diagrams). This is in line with our initial PCA (Figure 2G, H). Interestingly, PCA of common DEPs (Figure 1-source data 1, 2), i.e. DEPs in dependence on sex in both age groups, enabled effective discrimination between female and male samples, suggesting that these DEPs might represent sex-specific protein signatures in mouse paw skin and SCN (Figure 5C, D; Figure 5-figure supplement 1A, B). Among 87 common DEPs in skin (Figure 1-source data 1), we again identified a set of proteins with strong age dependency (Figure 5–figure supplement 1C). Remarkably, these candidates largely match the ones found in age-dependent comparisons of skin (Figure 3B). Most of these belong to Cluster 3 (Figure 2–figure supplement 1A), which is characterized by increased abundance specifically in male skin, and is associated with pathways like ‘muscle contraction’, ‘ion transport by P-type ATPases’, ‘glycolysis’, ‘metabolism of nucleotides and glycogen’. Overall, GO-BP analysis revealed that sex-dependent DEPs in skin and SCN are annotated to diverse pathways with only one pathway being shared in either age groups, i.e. ‘cornification’ in skin and ‘regulation of complement activation’ in SCN at 4 weeks and 14 weeks, respectively (Figure 5–figure supplement 2A, B). Moreover, we noticed that the complement activation pathway together with several processes of immune response were significantly enriched in both skin and SCN (Figure 5E, F), and more so in sex comparisons of 14 weeks tissues (green circles in Figure 5E, F). These biological processes are known to be affected under diverse pathological conditions (Ray *et al*., 2019; Wangzhou *et al*., 2021; Zeis *et al*., 2016). Thus, revealing their sexual dimorphism provides a crucial guide for adequate experimental design in future studies, especially when using rodent disease models.

**Figure 5:**
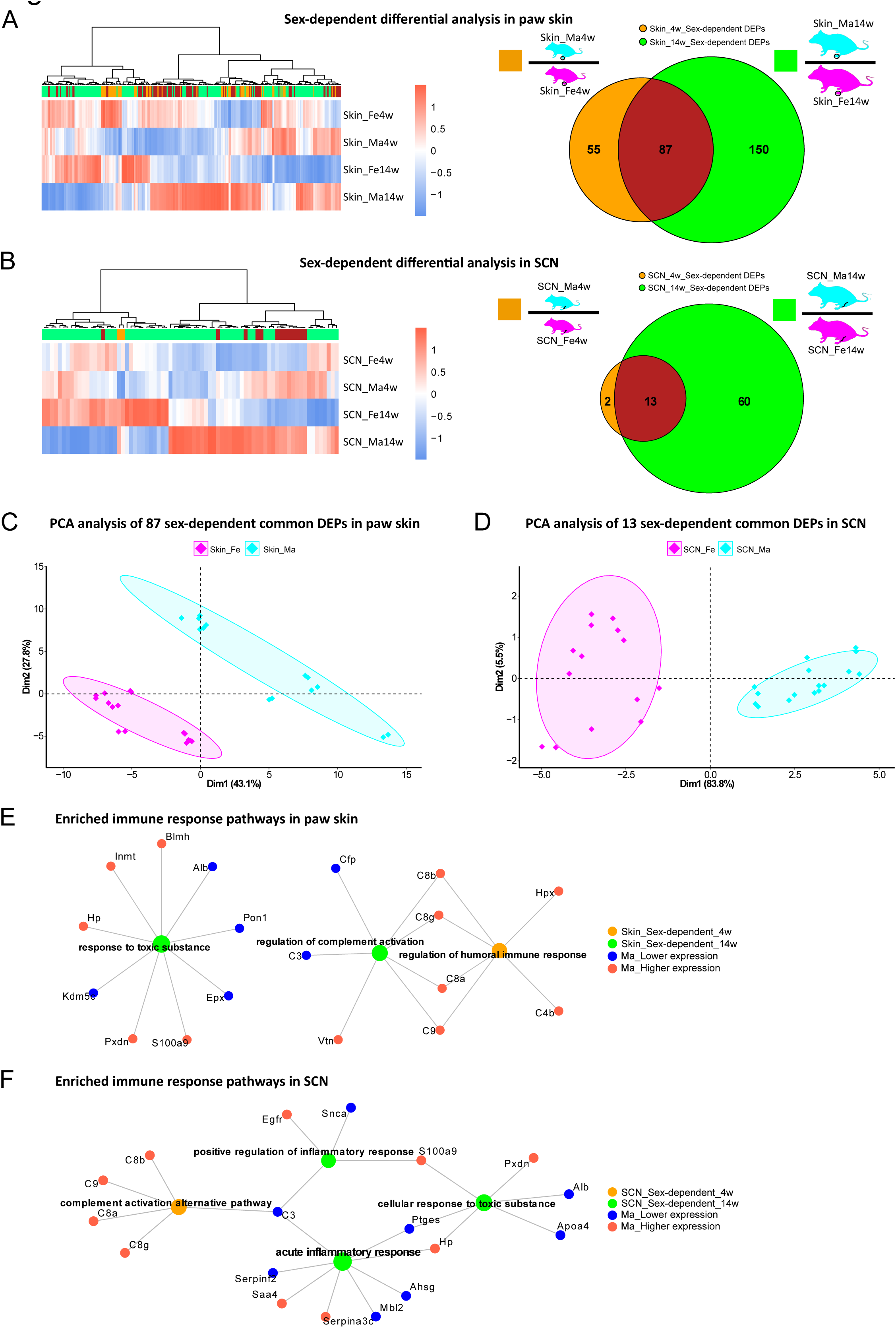
Differential expression analysis indicates protein signatures of sexual dimorphism in paw skin and SCN. A-B: DEPs of sex-dependent comparisons at 4 weeks and 14 weeks in paw skin (A) and SCN (B). Heatmaps show the normalized protein expression (averaged intensity) across age and sex groups. Venn diagram depicts sex-dependent DEPs at 4 weeks (yellow) and 14 weeks (green). C-D: Principal component analysis (PCA) reveals sex as an effective discriminator in paw skin and SCN tissues; females (magenta) and males (cyan). E-F: Visualization of enriched pathways related to immune response. Red: higher expression in males; blue: lower expression in males; green: pathways enriched at 14 weeks in a sex-dependent manner; yellow: pathways enriched at 4 weeks in a sex-dependent manner.

Next, we compared our datasets with previously published literature on molecular sex differences. Unfortunately, we did not find any previously published resource that assessed sex differences in mouse SCN. Therefore, we compared our SCN datasets with sex-associated genes found in human tibial nerve (Ray *et al*., 2019) (Figure 5-source data 1). However, most of the sex-dependent genes in human tibial nerves rather exhibited age- and not sex-dependent differences in our mouse SCN data, potentially due to species differences (Ray *et al*., 2019) (Figure 5–figure supplement 2C). A previous transcriptomics study on mouse hind paw skin revealed 123 differentially expressed genes (DEGs) comparing female with male mice aged of 8 to 12 weeks (Mecklenburg *et al*., 2020). Among these 123 DEGs, we quantified 43 proteins in our skin dataset, of which 11 were sexually dimorphic both at 4 weeks and 14 weeks (Figure 5-source data 2, Figure 5–figure supplement 2D) in line with published data (Mecklenburg *et al*., 2020). Reasons as to why we did not observe sex-dependent differences in the remaining 35 out of these 46 proteins could be manifold: starting with the broad age range used in the transcriptome study (Mecklenburg *et al*., 2020) to the known fact that transcript levels only show limited correspondence with protein expression levels (Maier *et al*., 2009). The latter highlights the importance to perform profiling studies on the proteome.

### Multiple proteins associated with skin diseases and pain exhibit age and sex dependency

In light of translational research, reverse translation of human data to mouse models of skin diseases is of high utility. Therefore, we compared our skin datasets with a list of top candidates (skin disease transcriptomic profiles, https://biohub.skinsciencefoundation.org/) found to be regulated in the skin of human patients suffering from diverse skin diseases such as psoriasis, acne, atopic dermatitis and rosaceae. Intriguingly, our proteome results harbor 347 out of 907 disease genes, of which 122 DEPs showed significant age and/or sex dependency (Figure 6A, Figure 6-source data 1). The latter was particularly prominent in disease candidates of psoriasis, atopic dermatitis, and rosaceae. Fuzzy C-means clustering analysis of these 347 skin disease-related proteins revealed not only discrete abundance patterns among age and sex groups but also proteins with differential profiles when comparing all 4 experimental groups (Figure 6-source data 2). For example, proteins in Cluster 2 exhibited low abundance in 14 weeks mice of both sexes, whereas proteins in Cluster 7 showed higher abundance only in 14 weeks male skin (Figure 6B). Discrete abundance patterns in dependence on age and sex were also observed on the level of individual DEPs (examples are given in Figure 6C; full list detailed in Figure 6-source data 1). Most proteins were less abundant at 14 weeks in both male and female samples despite showing significant sex differences, such as Collagen alpha-1(V) chain (Col5a1), Collagen alpha-2(V) chain (Col5a2), Fatty acid 2-hydroxylase (Fa2h), and Peroxidasin homolog (Pxdn). In contrast, Cellular retinoic acid-binding protein 2 (Crabp2) and Keratin, type II cytoskeletal 6A (Krt6a) represent examples of being more abundant at 14 weeks in both sexes. Others appear more highly expressed in 4 weeks female skin but show the opposite trend in male skin, e.g. Ras-related protein Rab-27A (Rab27a), Sushi repeat-containing protein SRPX2 (Srpx), and Versican core protein (Vcan). Overall, our data suggest significant regulation of human skin disease profiles by age and sex in mice – knowledge of utmost significance for reverse translational studies on the skin.

**Figure 6:**
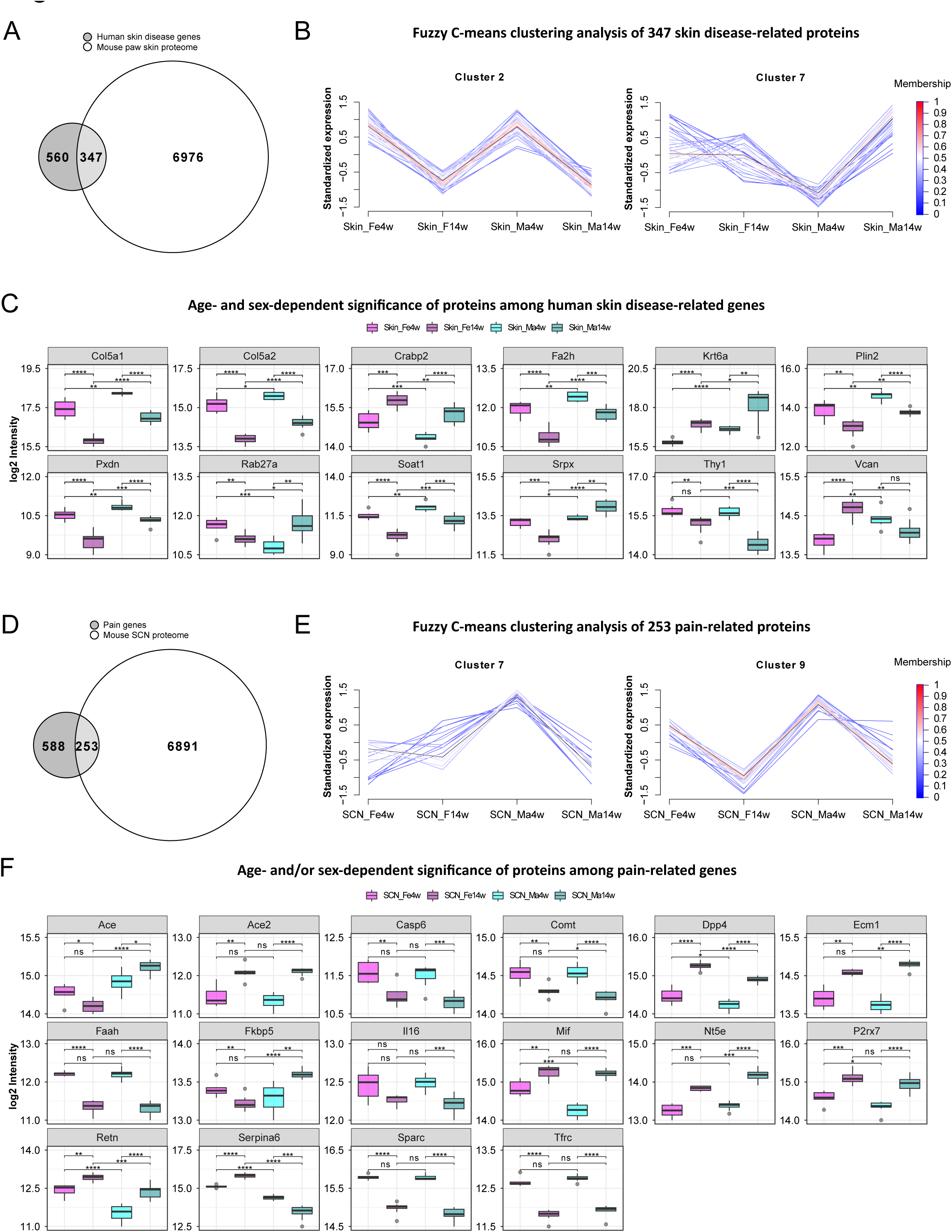
Multiple proteins associated with skin diseases and pain exhibit age and sex dependence. A: Venn diagram indicates the number of quantified protein IDs in paw skin (white) associated with various human skin diseases (light grey) upon comparison with a skin disease database (dark grey, https://biohub.skinsciencefoundation.org/). B: Examples of fuzzy C-means clustering analysis of the 347 protein IDs associated with human skin diseases illustrate their relative expression in experimental groups (other clusters are detailed in Figure 6-source data 2). C: Significantly expressed protein IDs associated with human skin diseases show both age and sex dependency. D: Venn diagram indicates the number of quantified protein IDs in SCN (white) associated with pain (light grey) upon comparison with known pain genes (dark grey). Pain-related genes are downloaded from publicly available pain gene databases: https://www.painresearchforum.org/, https://humanpaingeneticsdb.ca/hpgdb/, and http://paingeneticslab.ca/4105/06_02_pain_genetics_database.asp. E: Examples of fuzzy C-means clustering analysis of the 253 protein IDs associated with pain illustrates their relative expression in experimental groups (other clusters are detailed in Figure 6-source data 2). F: Significantly expressed protein IDs associated with pain show both age and sex dependency. Significance levels in C and F are indicated as: ns (q-value > 0.05), * (q-value ≤ 0.05), ** (q-value ≤ 0.01), *** (q-value ≤ 0.001), and **** (q-value ≤ 0.0001).

The SCN is affected by diverse pathologies such as nerve injury and neuropathic pain (Chen *et al*., 2021; Hildreth *et al*., 2009). In this context, we assessed the presence of known pain-related genes in SCN datasets by comparison with three publicly available pain gene databases (https://www.painresearchforum.org/, http://paingeneticslab.ca/4105/06_02_pain_genetics_database.asp, and https://humanpaingeneticsdb.ca/hpgdb/). Among 841 pain genes, 253 were quantified in our SCN proteome (Figure 6D, Figure 6-source data 3). Similar to skin disease-related proteins, fuzzy C-means clustering analysis revealed distinct abundance profiles for these 253 pain-related proteins differing by age and/or sex (Figure 6E, Figure 6-source data 2). Intriguingly, 111 of these pain-related proteins displayed significant changes (q-value ≤ 0.05, absolute log2 FC ≥ 0.38) by age and/or sex (Figure 6-source data 3), of which examples are given in Figure 6F. For instance, Angiotensin I converting enzyme (Ace) exhibits pronounced age and sex differences while its family member Ace2 only showed age-differences (i.e. higher expression in adult mice). Ace2 has been described to be associated with increased risk of non-specific orofacial symptoms in the OPPERA (Smith *et al*., 2013) prospective study and Ace is linked to migraine and potentially higher frequency as well as susceptibility (Dandona *et al*., 2007). Generally, Ace plays an essential role in vascular physiology and inflammation within the renin-angiotensin system (RAS). Unlike Ace2, catechol-O-methyltransferase (Comt) expression appears to be downregulated throughout maturation until adulthood in both sexes (Figure 6F). Comt variants assume diverse roles in the context of pain from its association with temporomandibular disorder (TMD) (Diatchenko *et al*., 2005) to postoperative pain (Belfer *et al*., 2015; Machoy-Mokrzyńska *et al*., 2019) and lower back pain (Omair *et al*., 2015). The FK506 binding protein 51 (Fkbp5) also displayed significant age and sex differences, with its expression being lower in female adults but higher in male adults compared to 4 weeks mice. Fkbp5 is a glucocorticoid receptor co-chaperone, and its polymorphisms predict persistent musculoskeletal pain after traumatic stress exposure (Bortsov *et al*., 2013; Linnstaedt *et al*., 2018). Moreover, it is generally involved in the acute stress response linked to diverse stress-related disorders in humans via the hypothalamic-pituitary-adrenal (HPA) axis (Häusl *et al*., 2021). Interestingly, variations in serpin peptidase inhibitor clade A member 6 (Serpina6), another pain gene with pronounced age differences and sex-opposite expression, are also associated with the HPA stress axis impacting the susceptibility to musculoskeletal pain (Holliday *et al*., 2010), the risk of cardiovascular disease as well as gene expression in peripheral tissues (Crawford *et al*., 2021). These few examples highlight the importance of considering changes in the abundance of aforementioned proteins across ages and in both sexes for research on diverse preclinical disease models. Therefore, our datasets represent a highly valuable and unique resource for biomedical studies. To further illustrate this utility, we have critically inspected our previous SCN datasets derived from the neuropathic pain model of nerve injury (spared nerve injury, SNI-model) in male adult mice (Barry *et al*., 2018). Indeed, several candidate proteins we previously reported (Barry *et al*., 2018) to be regulated upon SNI in adult males (i.e. Apoa1, C3, Postn, Pzp, Pcsk1n, Clu, Dpep1, Fabp3 and Egfr), exhibited age-dependent abundance differences in males with opposite trends in females (Figure 6-source data 4) – a fact, which has not been known before and might crucially affect SNI-induced pathology. Taken together, our data emphasize the essential need for male versus female as well as age-matched biomedical studies in parallel.

## Discussion

Age and sex as parameters in rodent-based research are known to strongly affect experimental outcomes *in vivo* and *in vitro*. Despite this knowledge we still do not know the molecular set-up of most cell and tissue types in dependence on age and sex (Garcia-Sifuentes *et al*., 2021; Woitowich *et al*., 2020). This gap is especially eminent at the proteome level – a fact that hampers our understanding of the molecular signature underlying physiological processes, diseases, and how they are impacted by age and sex as biological variables. Here, we present an optimized workflow of quantitative proteomics, which utilizes DIA-PASEF followed by data analysis with the publicly available program DIA-NN (Demichev *et al*., 2020). This optimized workflow enabled us to deeply profile and quantify the proteome of paw skin and sciatic nerve (SCN) in adolescent versus adult male and female naïve mice. Strikingly, our data reveal unprecedented insights into hitherto unknown age and sex differences of biological processes relevant for skin and SCN physiology as well as associated disorders. Therefore, our work fills an eminent gap defining the protein compendium of mouse skin and SCN and changes thereof in dependence on age and sex – serving as a unique resource for the scientific community. Even more, our work assertively highlights the significance of appropriate age and sex matching and provides a stepping stone for optimizing preclinical and translational research towards enhanced reproducibility and success.

We focused our analysis on paw skin and SCN isolated from 4 weeks and 14 weeks old male and female mice. Both tissue types are crucially implicated in diverse diseases. On one hand, in various allergic, itchy and inflammatory skin pathologies such as atopic dermatitis, psoriasis and lupus erythematodes. The SCN, on the other hand, is affected by a wide variety of motor and sensory neuropathologies induced by inflammation, trauma, and demyelination. Moreover, both the skin and SCN are involved in nociception and pain including chronic conditions. Our reasoning for the chosen age groups was as follows: given time and budget constraints as well as “comparability to historical data” (Reiber *et al*., 2022), it has become standard practice to perform experimental studies with mice of a wide age range, generally from 4 weeks to 12 weeks of age (Jackson *et al*., 2017; Reiber *et al*., 2022), regardless of the studied biological system. Of note, mice at 3-4 weeks of age are mostly used for cell culture-based *in vitro* studies (Boyer *et al*., 1994; Isensee *et al*., 2017; Lin *et al*., 2018; Malin *et al*., 2007). For example, peripheral sensory neurons of DRG exhibit better health and growth factor-dependent survival when isolated from young rodents (Malin *et al*., 2007; Melli *et al*., 2009). Similarly, Schwann cells originated from younger human donors proliferated faster than those from older donors (Boyer *et al*., 1994; Monje, 2020). In contrast to these younger ages used for cell culture-based research, most studies on mouse behavior are conducted at ages of 6-12 weeks, i.e., a period with overt maturational changes before reaching adulthood from 12 weeks of age onwards (Flurkey *et al*., 2007). Notable examples of this practice include studies on cutaneous touch, somatosensation and (chronic) pain, and diseases of the central and peripheral nervous system just to name a few (Narayanan *et al*., 2016; Poole *et al*., 2014; Zheng *et al*., 2019). Furthermore, the most extensive and highly valuable RNA-seq-based resource describing the molecular set-up of mouse CNS (central nervous system) and PNS (peripheral nervous system) cell types (Zeisel *et al*., 2018) (https://www.mousebrain.org) was assembled by pooling mice of both sexes aged 2-3 weeks, as well as 6 and 8 weeks. Our results show that the proteome exhibits clear differences when comparing 4 weeks with 14 weeks mice raising concerns about aforementioned practices of comparing and correlating *in vitro* data derived from young ages with *in vivo* data of older ages. Failed correlations among these age groups may have led to false negatives and prevented new findings. On the contrary, mechanisms discovered in young cells may be wrongly accounted for phenotypic differences observed in adult rodents. Based on our data, we strongly suggest suitable age matching across different methods within a study to ensure scientific rigor and reproducibility.

From a technical point of view, we have here employed a highly sensitive workflow based on DIA-PASEF followed by data analysis via DIA-NN (Demichev *et al*., 2020). This enabled us to provide the most in-depth and highly reproducible proteome dataset of mouse skin and SCN published thus far. While we have previously used DIA-MS, but not DIA-PASEF, to investigate various tissues (Barry *et al*., 2018; Rouwette *et al*., 2016; Sondermann *et al*., 2019) in mouse pain models, we have not achieved this high quantitative depth as reported here. For instance, in SCN, we quantified more than double of proteins (8400 IDs, Figure 2C) compared to our previous study using DIA-MS (3473 IDs) (Barry *et al*., 2018). DIA workflows generally facilitate higher reproducibility of quantitative profiling (Domon *et al*., 2010), however, resulting MS spectra are very complex requiring careful interpretation (Bilbao *et al*., 2015). To address that, improvements of diverse aspects have been continuously implemented. For instance, from the hardware point of view, the addition of ion mobility separation to the chromatographic and mass separation of peptides has significantly reduced spectral complexity (Helm *et al*., 2014) in DIA-MS. Furthermore, the DIA-PASEF workflow was developed on timsTOF Pro instruments (Bruker Daltonik), which enables nearly complete sampling of the precursor ion beam (Meier *et al*., 2020). On the other hand, new algorithms have been developed to interpret complex spectra. For example, Arnaud Droit and colleagues systematically evaluated and compared the most used DIA-data processing software (Gotti *et al*., 2021) (DIA-NN, DIA-Umpire, OpenSWATH, ScaffoldDIA, Skyline, and Spectronaut): DIA-NN outperformed others in terms of peptide and protein identifications (Demichev *et al*., 2020). Given that DIA-PASEF has not been widely used yet, above all not for mouse skin and SCN, we have systematically compared the more commonly used DDA-PASEF mode with DIA-PASEF. We specifically compared (i) the number of protein IDs quantified, (ii) the coefficients of variation (CVs), and (iii) quantitative correlations between technical replicates of each sample. As expected, DIA-PASEF outperformed DDA-PASEF in all parameters (Figure 1). Consequently, we obtained all here reported datasets on paw skin and SCN via DIA-PASEF coupled with DIA-NN library-free data analysis.

Both tissues, however, harbor diverse cell types such as keratinocytes, immune cells, peripheral nerve endings, and fibroblasts in skin and glia cells, immune cells and axons in SCN. Because of this cellular complexity, we cannot assign the detected age-and sex-dependent proteome differences to specific cell types – a limitation applicable to all -omics assays, if not performed on the single-cell level. In contrast to established procedures for single-cell RNA-seq, proteomics on single cells is still in its infancy. Yet, the transcriptome cannot predict disease phenotypes in a straight forward manner given the highly dynamic regulation of proteins by cellular buffering mechanisms regulating their synthesis and turnover (Liu *et al*., 2016; Schwanhäusser *et al*., 2011). Consequently, pathologies and associated changes in molecular and cellular signaling need to be monitored on the proteome level. This calls for the development of technological advances for accurate, highly sensitive, and comprehensive proteome profiling on the single-cell level. Recently, Brunner *et al*. established an ultra-high sensitivity MS workflow to quantify a single-cell proteome from HeLa cell culture with protein IDs ranging from 1018 (cell cycle G1 phase) to 1932 (G1/S phase) per single cell (Brunner *et al*., 2022). Though this pipeline has provided the opportunity to analyze single-cell derived proteomes, the workflow requires elaborate sample preparation and technical equipment. Furthermore, it has not yet been applied to complex tissues harboring multiple cell types.

In conclusion, here, we present the most extensive proteome compendium of mouse skin and SCN described thus far. Our work demonstrates prominent and previously unknown sexual and age dimorphisms in paw skin and SCN of naïve mice. Many of the here reported differences are likely relevant for our mechanistic understanding of various disorders involving the skin (e.g. inflammatory pathologies like atopic dermatitis and psoriasis) and SCN (motor- and sensory neuropathologies induced by inflammation, trauma, and demyelination). Therefore, our study serves as a unique resource for different life science disciplines. Even more, our work advocates for the importance of appropriate age and sex matching and provides new avenues for improving the reproducibility, generalizability, and success of preclinical and translational research on skin and SCN.

## Data availability

All datasets included in this study are listed in 14 source data files. Moreover, proteome raw data generated in this study were deposited to the PRIDE archive via ProteomeXchange (https://www.proteomexchange.org) with identifier: PXD034476 (Username for reviewers, Password: GqgXFQHX).

## Ethics

All mouse work strictly followed the regulations stated in the German animal welfare law (TierSchG §4). For this study, mice were sacrificed by CO_2_ to obtain tissues, and no other procedure was performed. Therefore, mouse work of this study is not considered an animal experiment according to §7 Abs. 2 Satz 3 TierSchG. All procedures were approved and supervised by the animal welfare officer and the animal welfare committee of the Max Planck Institute for Multidisciplinary Sciences, Göttingen, Germany. The animal facility at the Max Planck Institute for Multidisciplinary Sciences is registered according to §11 Abs. 1 TierSchG.

## Author contributions

Conceptualization: FX, JRS, DGV & MS; Methodology: FX & JRS; Validation: FX, JRS, DGV & MS; Formal analysis: FX & JRS; Investigation: FX & JRS; Data curation: FX, JRS & MS; Writing original draft: FX, JRS & MS; Writing, review & editing: FX, JRS, DGV & MS; Visualization: FX; Supervision: MS; Project administration: FX, JRS, DGV & MS; Funding acquisition: MS.

## Acknowledgement

We would like to thank Tanja Nilsson (Max Planck Institute for Multidisciplinary Sciences, Göttingen, Germany) and Sabrina Grundtner (Division of Pharmacology & Toxicology, University of Vienna, Austria) for their efforts regarding tissue isolation. We are grateful to the Systems Biology of Pain team at University of Vienna for discussions, and to Daniel Segelcke (University Hospital Muenster, Muenster, Germany) for critical reading of the manuscript. We also appreciate the help from the Bruker Daltonik team on setting up the proteomics platform.

